# Evolutionary determinants of morphological polymorphism in colonial animals

**DOI:** 10.1101/046409

**Authors:** Carl Simpson, Jeremy B.C. Jackson, Amalia Herrera-Cubilla

**Affiliations:** Department of Paleobiology, National Museum of Natural History, Smithsonian Institution, Washington, D.C. 20013-7012, USA; Smithsonian Tropical Research Institute, 0843-03092, Balboa, Republic of Panama

**Keywords:** Coloniality, polymorphism, division of labor, life-history, Bryozoans

## Abstract

Colonial animals commonly exhibit morphologically polymorphic modular units that are phenotypically distinct and specialize in specific functional tasks. But how and why these polymorphic modules have evolved is poorly understood. Across colonial invertebrates there is wide variation in the degree of polymorphism from none in colonial ascidians, to extreme polymorphism in siphonophores such as the Portuguese Man of War. Bryozoa are a phylum of exclusively colonial invertebrates that uniquely exhibit almost the entire range of polymorphism from monomorphic species to others that rival siphonophores in their polymorphic complexity. Previous approaches to understanding the evolution of polymorphism have been based upon analyses of (1) the functional role of polymorphs or (2) presumed evolutionary costs and benefits based upon evolutionary theory that postulates polymorphism should only be evolutionarily sustainable in more “stable” environments because polymorphism commonly leads to the loss of feeding and sexual competence. Here we use bryozoans from opposite shores of the Isthmus of Panama to revisit the environmental hypothesis by comparison of faunas from distinct oceanographic provinces that differ greatly in environmental variability and we then examine the correlations between the extent of polymorphism in relation to patterns of ecological succession and variation in life histories. We find no support for the environmental hypothesis. Distributions of the incidence of polymorphism in the oceanographically unstable Eastern Pacific are indistinguishable from those in the more stable Caribbean. In contrast, the temporal position of species in a successional sequence is collinear with the degree of polymorphism because species with fewer types of polymorphs are competitively replaced by species with higher numbers of polymorphs on the same substrata. Competitively dominant species also exhibit patterns of growth that increase their competitive ability. The association between degrees of polymorphism and variations in life histories is fundamental to understanding of the macroevolution of polymorphism.

## Introduction

Modules of colonial organisms originate asexually as clones and typically remain physically and physiologically connected (Beklemishev 1969; Jackson 1979a). Colonies reproduce sexually although clonal reproduction of new colonies is common (Hughes and Cancino 1986; Jackson 1986; Jackson and Coates 1986). Some, but not all, colonial organisms evolve phenotypic variation among their clonal members that ranges from simple shape differences of component modules (Cheetham 1973) to profoundly disparate polymorphic forms (McKinney and Jackson 1991; Ryland 1970). Bryozoan species exhibit a remarkable range of modular polymorphism, from uniform colonies within which every member is identical to colonies so polymorphic they rival even siphonophores in the large numbers of distinct modular types they contain. In highly polymorphic bryozoans, phenotypic differences among modules are accompanied by the functional loss of many organ systems, including feeding and sexual competency (Ryland 1970; Carter et al. 2010; Ostrovsky 2013). Darwin, in the sixth edition of the Origin (Darwin 1872), recognized the difficulty inherent in understanding the evolutionary differentiation of bryozoan polymorphs. Even now, there is no definitive understanding of the problem.

The evolutionary problem of polymorphism is related to the evolution of sex in that there is an inevitable cost of differentiation caused by loss of feeding and sexual competency. These costs must be recouped by the particular adaptive benefit associated with differentiating polymorphs. Any general understanding of the evolution of polymorphism must take this cost and its balancing benefits into account.

Ever since Darwin, scientists have explored two explanations for the evolution of polymorphism. The first, similar to that advocated by Darwin, tries to understand the evolution of different types of polymorphs by understanding the functions each type performs in a colony. Unfortunately, except in rare cases such as the worm and leg pinching ability of the bird’s-head avicularia of *Bugula* (Ryland 1970), the functions of bryozoan polymorphic types are too poorly understood for meaningful analysis. The second explanation for the evolution of polymorphism is as a response to the stability of the external environment (Hughes and Jackson 1990; Schopf 1973). Unstable environments limit the number of different polymorph types and more stable environments relax those limits. For bryozoans, this way of understanding was inspired by the ergonomic theory of caste differentiation in social insects (Oster and Wilson 1979).

Neither approach has yielded a general understanding of the problem of modular polymorphism. Instead we have only incomplete snippets of functional information and phenomenological descriptions. We addressed this problem in two ways. First, we revisit the environmental hypothesis at an intermediate oceanographic spatial scale more appropriate to directly testing the hypothesis. Second, we investigated the functional hypothesis by exploring correlations between polymorphism and patterns of ecological succession and variation in life histories. Patterns of succession provide good summaries of the adaptive benefits of polymorphism because they are the result of successful adaptive strategies. Furthermore, life-history strategies show, in a broad sense, how the fitness benefits of those adaptive strategies are achieved.

## The nature of bryozoan polymorphism

Bryozoans are colonial animals that grow by budding modules termed zooids, each of which is homologous to a single solitary animal such as an ant or snail (McKinney and Jackson 1991; Ryland 1970). Colonies are tessellated by generic feeding zooids termed autozooids. Scattered among the autozooids are polymorphic zooids that are commonly discrete in shape and function (Fig. 1). Autozooids feed by protrusion of a tentacular polypide animal through the orifice of the zooid that is protected by a hinged flap called an operculum. Autozooids can also produce gametes as well as perform all of the basic tasks required for survival, and are considered the evolutionarily primitive condition (Banta 1973; Cheetham 1973; Cheetham et al. 2006; Silén 1977). Polymorphic zooids within a single colony may differ more from each other than even the most extreme members of discrete social insect castes (Wilson 1975). Moreover, the number and frequency of different types of polymorphs can vary from colony to colony both within and among species.

**Figure 1:**
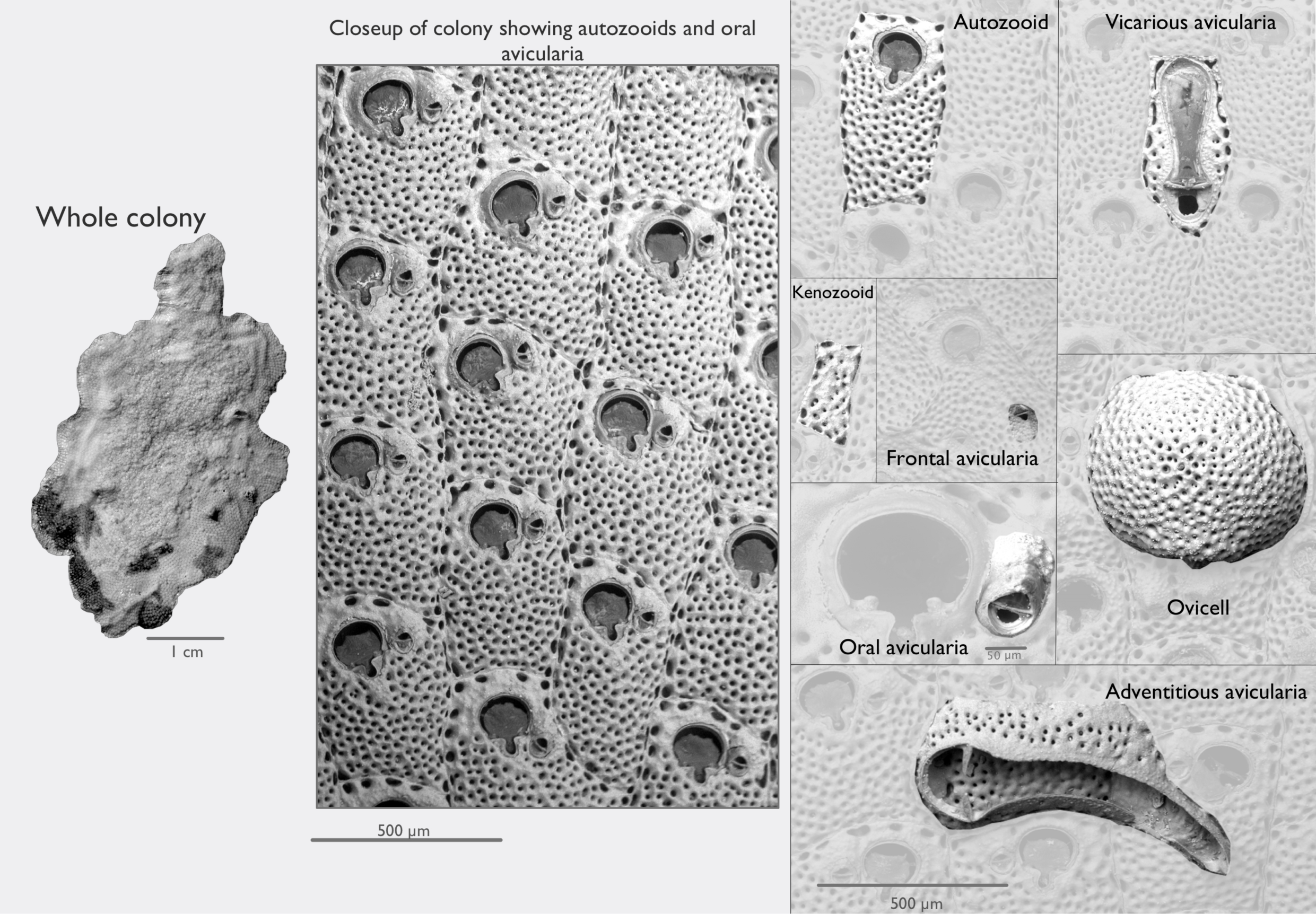
The zooids that constitute a bryozoan colony can be highly polymorphic. Whole colonies can be many tens to hundreds of square centimeters in area, but the thousands of zooids that make the colony are small, most commonly less than a millimeter in length. Zooids in encrusting species like this *Stylopoma* new species 3 (Jackson and Cheetham 1994) form a mosaic across their substratum. As the colonies grow, they radiate out from the oldest part of the colony, which is often near the center, as seen in the left image of a whole *Stylopoma* colony. Zooming into the colony shows zooid-level detail. Each repeated structure in the central image consists of two animals, an autozooid with an oral adventitious avicularium. *Stylopoma* autozooids protrude their tentacle crowns through a hatched orifice that is about 100 *¼*m wide. Scattered across this colony (right panel) are six polymorph types. This colony has a single type of vicarious avicularium that replaces an autozooid inline. There is a type of kenozooid, which fills space. This species has ovicells, which commonly occur in a band halfway between the edge and the center of the colony, and three types of adventitious avicularia: an oral, frontal, and a large type that overgrows many zooids.

The various types of polymorphs differ from autozooids by giving up one or more basic tasks to specialize in particular functions. Avicularia are the most common polymorphs and come in many forms and sizes, but they all share some basic attributes. Their hinged operculum is modified so that, like a person waving a flag, it can be moved about across the surface of the colony. Avicularia reduce or lose other body parts, such as the polypide with its tentacle crown and major organs, though a vestigial polypide can provide a sensory function within many types of avicularia (Carter et al. 2010). Because avicularia are so reduced, they cannot feed or contribute to reproduction, so they remain connected to other zooids and depend on them for nutrition. The tasks that avicularia perform in the colony include cleaning, defense, hygiene, and patterning the flow of water across the colony surface (Carter 2008; Winston 1984; Winston 2010), although the specific functions of the various types of avicularia are poorly understood. Vicarious avicularia (Fig. 1) are budded with other zooids in the main colony plane, replacing an autozooid without changing the mosaic pattern. Adventitious avicularia (Fig. 1) are budded atop the plane of the colony where they may overgrow preexisting zooids, adding a second level. Adventitious avicularia are commonly found adjacent to the orifice of autozooids (Fig. 1), scattered across their frontal walls (Fig. 1), or encrusting other parts of the colony like salt on a pretzel (Fig. 1).

There are three other major types of polymorphs: spines, kenozooids, and ovicells (Fig. 1). Spines, also known as spinozooids (Silén 1977), are zooids that have been reduced into small spiny skeletal protrusions. Nevertheless, each spine is separated from the adjacent autozooid by cuticular tissue (Silén 1977), just as autozooids themselves are separated from each other (Ryland 1970). Kenozooids are diminutive zooids that fill gaps in the colony’s mosaic where a full autozooid or avicularium would be too large. Although it may seem trivial, filling space is crucial for protection from competitive overgrowth by other organisms that have the potential to get a foothold in empty areas (Jackson 1979b; McKinney and Jackson 1991; Palumbi and Jackson 1982).

Ovicells are dome-shaped polymorphs whose sole function is to brood larvae (Fig. 1). Ovicells are budded from the distal end of a parental autozooid, placing the orifice of an ovicell within reach of the parental autozooid polypide’s tentacle crown. Ovicells vary among species in their position relative to the surface of the colony. In some species they remain submerged below the colony surface within the parental autozooid, but in others the ovicells stand in stark relief above the frontal surfaces of nearby autozooids. Unfertilized eggs or embryos (the site of fertilization is unknown) are passed from the tentacles of the adjacent zooid to the ovicell where they are brooded until ready to disperse (Ostrovsky 2013). Ovicells are uncommon and are usually one or more orders of magnitude less frequent than autozooids within a colony (Simpson 2012).

## Two Hypotheses

Polymorphic differences among constituent animals (or modules of a colony) are often functional. Each distinct animal form within a society or a colony divides up the labor by specializing on a specific task, such as feeding, defense, or reproduction (Beklemishev 1969; Harvell 1994; Lidgard et al. 2012; Mackie 1986; Wilson 1975; Winston 1984). The division of labor among specialist forms is thought to dramatically increase efficiency over generalist forms, in which individual members are capable of doing everything alone (Beklemishev 1969; Huxley 1912). However, the efficiency gained by evolving polymorphism can be structured over macroevolutionary time in two ways. It can be structured environmentally or functionally. These two hypotheses roughly map onto how the distribution of species of particular degrees of polymorphism tend to be spatially structured. With the environmental hypothesis species with similar degrees of polymorphism will track similar environments. With the functional hypothesis, species with different degrees of polymorphism can co-occur as long as they differ in adaptive strategy.

According to the environmental hypothesis, polymorphism is both costly and beneficial. The costs come directly from reduction in the numbers of individuals or modules involved in feeding or sexual reproduction. Polymorphism should increase only if the colonies or societies persist long enough for the evolutionary benefits of specialization to outweigh the costs of decreased feeding or reproduction overall. Average longevity should in turn increase proportionally with increased environmental stability. Conversely, colonies living in unstable environments will, on average, die before the benefit of polymorphism is accrued. Therefore, the environmental hypothesis predicts that highly polymorphic colonies will be absent from unstable environments.

This hypothesis is also known as the ergonomic hypothesis because the number of polymorph types evolves to fit the stability of the environment (Schopf 1973; Oster and Wilson 1979). In social insects, the ergonomic hypothesis has largely fallen out of favor as the cause of caste type diversity (Bourke 2011). However, in sessile clonal organisms, the environmental hypothesis takes on added strength because success or death is directly linked to the stability of the substrates they grow upon and prefer (Jackson 1977; Jackson 1979b). Thus the environmental hypothesis has the potential to be a valid for sessile colonial organisms even though it is not supported in social insects. To evaluate this we examined the occurrence of polymorphism in species occurring in the strikingly different oceanographic regimes on opposite sides of the Isthmus of Panama.

Alternatively, the benefits of polymorphism may accrue through increased efficiency associated with the proliferation of specialized functions. To evaluate this, we observed how species with varying degrees of polymorphism sort themselves out along a successional sequence. Although an indirect measure of the payoff of possessing polymorphs, this approach provides a direct measure of the presence of those benefits exceeding the costs.

## Methods

Environmental stability varies over a wide array of spatial and temporal scales. Stability can range from very local—such as the ephemeral stability of a shell on the seafloor compared to a large boulder or coral—to larger scale differences in prevailing climactic or oceanographic conditions, such as the broad patterns of hurricane frequency across the tropics or along the eastern and western borders of continents (Jackson 1991; McKinney and Jackson 1991; Jackson and D’Croz 1997). All scales of environmental stability are important for the environmental hypothesis for polymorphism because the costs and benefits of polymorphism need to be balanced locally, within an individual colony, and also at larger scales as polymorphism evolves within a population and among species. At the smallest scale, the maximum lifetime of a sessile colony cannot exceed that of its substrate, as it cannot move. Similarly, at the largest scale, populations can persist only as long as their environments persist, unless they can tolerate a highly variable environment—in which case the costs and benefits of polymorphism must be accounted for in every environmental state. The costs in each state are important, because they are dominated by maintaining non-feeding and non-reproducing members. Moreover, costs and benefits in each environment may be at odds with each other, making an optimized strategy more difficult to evolve.

The environmental hypothesis has been tested in bryozoans twice at the two extremes of spatial scale. Schopf (1973) compared the incidence of polymorphism on a global scale between shallow, supposedly unstable tropical seas versus supposedly more stable, high-latitude and deep-sea environments. However, Schopf’s assumptions about oceanographic stability were incorrect, since highly stable and unstable environments exist in all three regions. In the second test, Hughes and Jackson (1990) compared levels of polymorphism along a narrow stretch of the Caribbean coast of Panama in relation to the size and physical stability of different sized substrata. They compared substrates ranging from small shells and pebbles that are easily buried and rolled about on the sea floor, up to large, firmly attached and long-lived corals and reef framework. They found no difference in the number of types of polymorphs in relation to the substrate stability at this small spatial scale.

Nevertheless, the environmental hypothesis remains untested by comparison of the levels of polymorphism from different oceanographic regimes. We therefore compared the incidence of polymorphism between the strikingly different shallow-water coastal environments on opposite sides of the Isthmus of Panama (D’Croz and O’Dea 2007; Jackson and D’Croz 1997; Lessios 2008; O’Dea and Jackson 2009; O’Dea et al. 2007). Eastern Pacific coastal environments exhibit strong seasonal upwelling with large fluctuations in temperature and planktonic productivity, as well as considerable interannual variation in oceanographic conditions that directly impact marine faunas (Baker et al. 2004; Colgan 1990; Glynn and Colgan 1992; Glynn et al. 2001). Development of coral reefs is meager and seagrass beds are absent. In contrast, Caribbean coastal environments exhibit low seasonality and planktonic productivity, and coral reefs and seagrass beds are extensive (Jackson and D’Croz 1997; O’Dea et al. 2007).

These environmental differences arose over the past 5 million years as the shallow seaways connecting the two oceans gradually closed, resulting in considerable evolutionary divergence in faunas (Lessios 2008; O’Dea et al. 2007; O’Dea et al. 2016). Consequently, bryozoans on either side of the Isthmus have had sufficient time to evolve a largely independent set of species, which have evolutionarily divergent differences in many aspects of their morphology (Jackson and Herrera-Cubilla 2000). Thus, if the environmental hypothesis applies at this intermediate scale, species of Caribbean bryozoans should have more types of polymorphs and the frequency distribution of the number of types of polymorphs should be shifted upward in the Caribbean relative to the Eastern Pacific.

Settling experiments using artificial substrates placed in the environment over varying lengths of time have been used to observe the successional sequence of species occurrence in the development of bryozoan communities (McKinney and Jackson 1991; Winston and Jackson 1984; Barnes and Sanderson 2000). We exploited Winston and Jackson’s results to record the number of different types of polymorphic zooids characteristic of the bryozoan species observed to be growing on these panels. Two sets of six replicate 15x15 cm square panels were set out in shallow water off of Jamaica in 1977. Each plate was observed 7 times over 3 years to map the occurrence, position, and percent cover of each of the ten species that varyingly settled and grew on the panels. We transformed the percentage cover data to calculate proportional cover over the course of the experiment for each species. We calculated the average the number of types of polymorphs across species by weighting each species by its relative proportional cover.

## Data

We estimated the total diversity of polymorphs among 79 species of ascophoran cheilostome bryozoans in Smithsonian collections obtained from shallow-water coastal environments on either side of the Isthmus of Panama (Jackson and Herrera-Cubilla 2000) (Table 1). All species are encrusting and are found on many substrates, from shells and pebbles, to coral rubble and reef-framework. Not all colonies express or preserve all of the polymorphs they are capable of producing, so whenever possible we tallied the mean and maximum number of polymorphic types observed from 1-28 colonies per species. We have counts for 40 species from 25 localities in the Eastern Pacific and 39 species from 10 localities in the Caribbean.

**Table 1:**
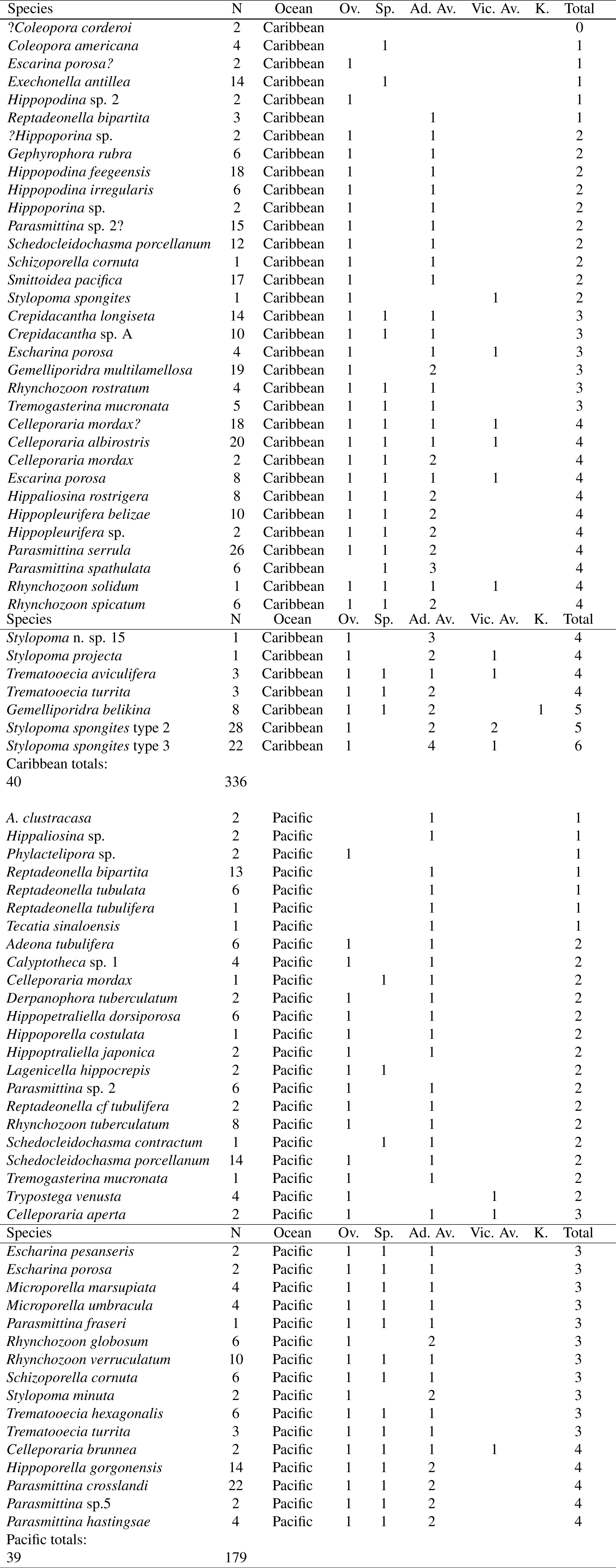
Polymorph types for ascophoran cheilostome bryozoan species from shallow coastal waters of the East Pacific and Caribbean of Panama. We show the number of distinct polymorphs of each type observed in a species, which maybe more than the maximum observed in any single colony. N = number of sampled colonies; Ov. = Ovicell; Sp. = spines; Ad. Av. = Adventitious Avicularia; Vic. Av. = Vicarious Avicularia; K. = kenozooid.

We recognized eight basic types of polymorphs that differ qualitatively in body plan and in the direction of budding from parent zooids (Fig. 1). These include: ovicells, kenozooids, spines, vicarious avicularia that occur inline with autozooids, and adventitious avicularia (which are frontally budded and can vary in shape and position). At least four different types of adventitious avicularia can be identified by their position on the zooidal or colony surface. Oral adventitious avicularia occur adjacent to the autozooid’s orifice. A second type of adventitious avicularium occurs on the zooidal frontal wall. A third type occurs on the ovicell in some species. The fourth type includes large adventitious avicularia that overgrow multiple zooids.

The total number of possible types of polymorphs exceeds the number actually observed in any species. Moreover, the eight polymorph types that we recognize here are a subset of the total number of possible polymorphs that occur in cheilostome bryozoans as a whole (Silén 1977). We lumped other named polymorph types together into one of the eight categories. However, we distinguished between polymorphs beyond our eight named types if they differed in shape from other polymorphs in the same positional category. For example, we considered vibriculae to be a bristly type of avicularium. Thus, if a species expressed both bristly and flap-like avicularia, we would count these as two different types of polymorphs.

## Results

The frequency distributions of numbers of types of polymorphs per species are statistically indistinguishable between the Eastern Pacific and the Caribbean (Fig. 2). An average bryozoan species in both the Eastern Pacific and the Caribbean has an ovicell, spines or a kenozooid, and one type of avicularium, or perhaps an ovicell and two types of avicularia. Eleven genera occur on both sides of the isthmus, yet the distributions of their polymorphs are not statistically distinct between the oceans based on a comparison of the maximum incidence of polymorphic zooid types (Wilcoxon W = 49, P = 0.45). The mean of the maximum observed number of polymorphs observed in each species is 2.79 in the eastern Pacific and 2.38 in the Caribbean (Wilcoxon W = 1039.5, P = 0.12). The median number of polymorphs for species is 2 in the Pacific and 3 in the Caribbean. If we incorporate the variation in expressed polymorphism from colony to colony within species by calculating the mean observed, we still find no difference between distributions (average for Eastern Pacific species = 2.2; average for Caribbean species = 2.03; Wilcoxon W = 961, P = 0.48). Our results are consistent with the pattern Hughes and Jackson (1990) found at much smaller spatial scales. The incidence of polymorphism in ascophoran bryozoans is not optimized ergonomically between the strikingly different coastal environmental conditions on opposite sides of the Isthmus of Panama.

**Figure 2:**
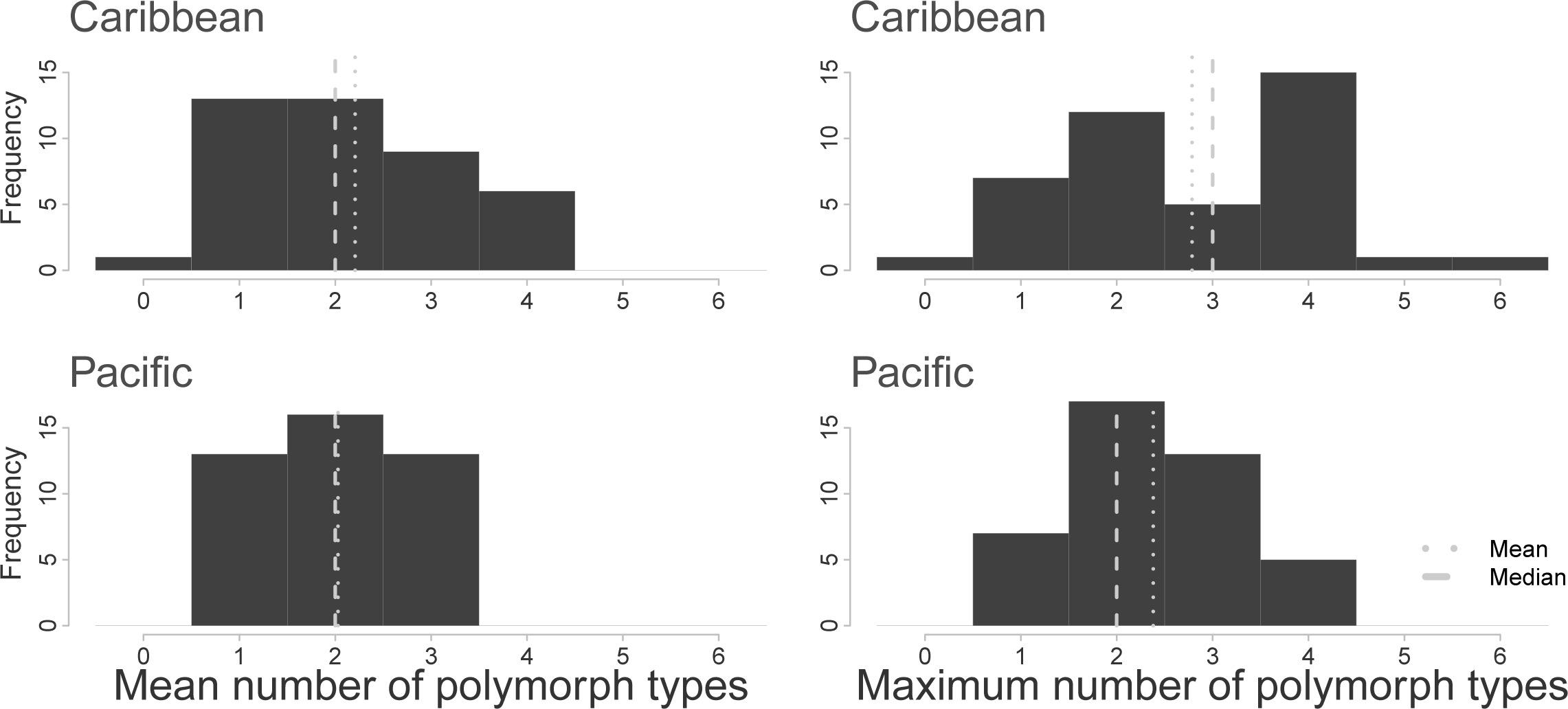
The frequency distributions of maximum and mean numbers of polymorph types observed in colonies of each species are compared between the shallow-water coastal bryozoans in the Caribbean and the East Pacific. We took the maximum and the mean numbers of polymorph types for 1-28 colonies per species. We compare 39 species in the Eastern Pacific to 40 species in the Caribbean.

Nevertheless, there are important differences between the oceans in the extremes of the distributions of polymorphism (Fig. 2). For example, there are 17 Caribbean species with 4 or more types of polymorphs versus only five such species in the Eastern Pacific. We find strong statistical support for an excess of highly polymorphic species in the Caribbean relative to the Eastern Pacific (*χ*^2^ = 7.45, df = 1, P = 0.006). Moreover, these highly polymorphic species increasingly dominated the fouling panels as ecological succession occurred over three years of the experiment (Fig. 3, Fig. 4; Spearman’s *ρ* = 0.64, P = 0.014).

**Figure 3:**
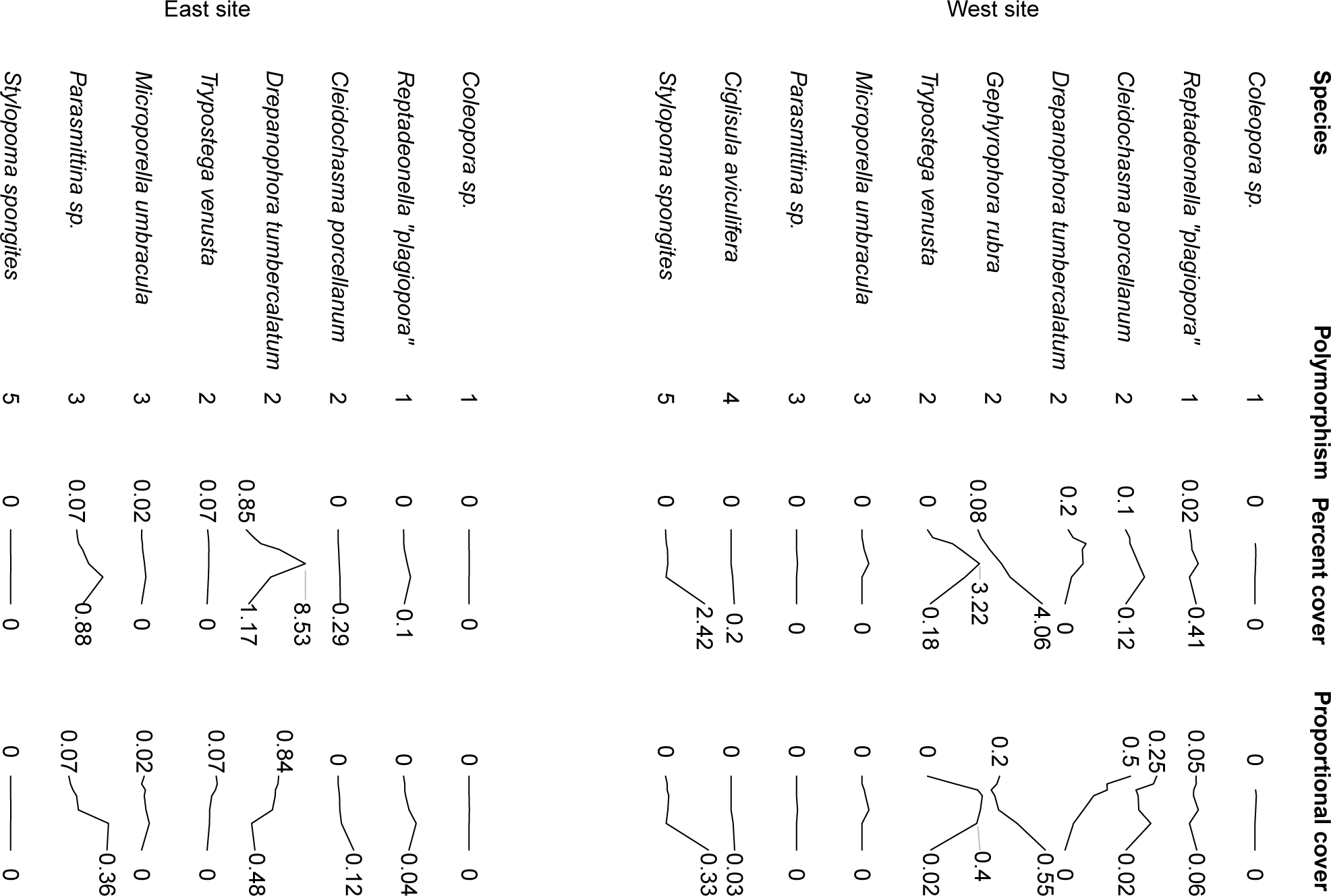
We used the results of a settling experiment that tracked the recruitment, growth, and mortality of encrusting bryozoans onto clean “fouling” panels (data from Table 2 in Winston and Jackson 1984). Two sets of six 15 cm square panels were set out approximately 100 m apart in 12-13 m water depth along the fringing coral reef on the west side of the embayment at Rio Brueno, Jamaica. A census was conducted for each plate 7 times over 3 years. During each census, 10 species of ascophoran bryozoans where evaluated for the percentage of the plate that their colonies covered. This set of 10 species happens to be a subset of the ones we surveyed in Table 1 and we pulled the observed degree of polymorphism from our Table 1. Here we show the incidence of polymorphism for each species, the percentage of the fouling plate covered by each species, and the proportional cover to show the relative dominance of each species over time. The percent and proportional cover are shown as sparkline timeseries that span the three year experiment. Each time series is scaled to others in its site and row. Start and ending percentages are shown and if the time series is strongly peaked the maximum percentage is also shown. The percent cover is the average replicate samples and are highly variable. The standard deviations can be found Table 2 in Winston and Jackson (1984).

**Figure 4:**
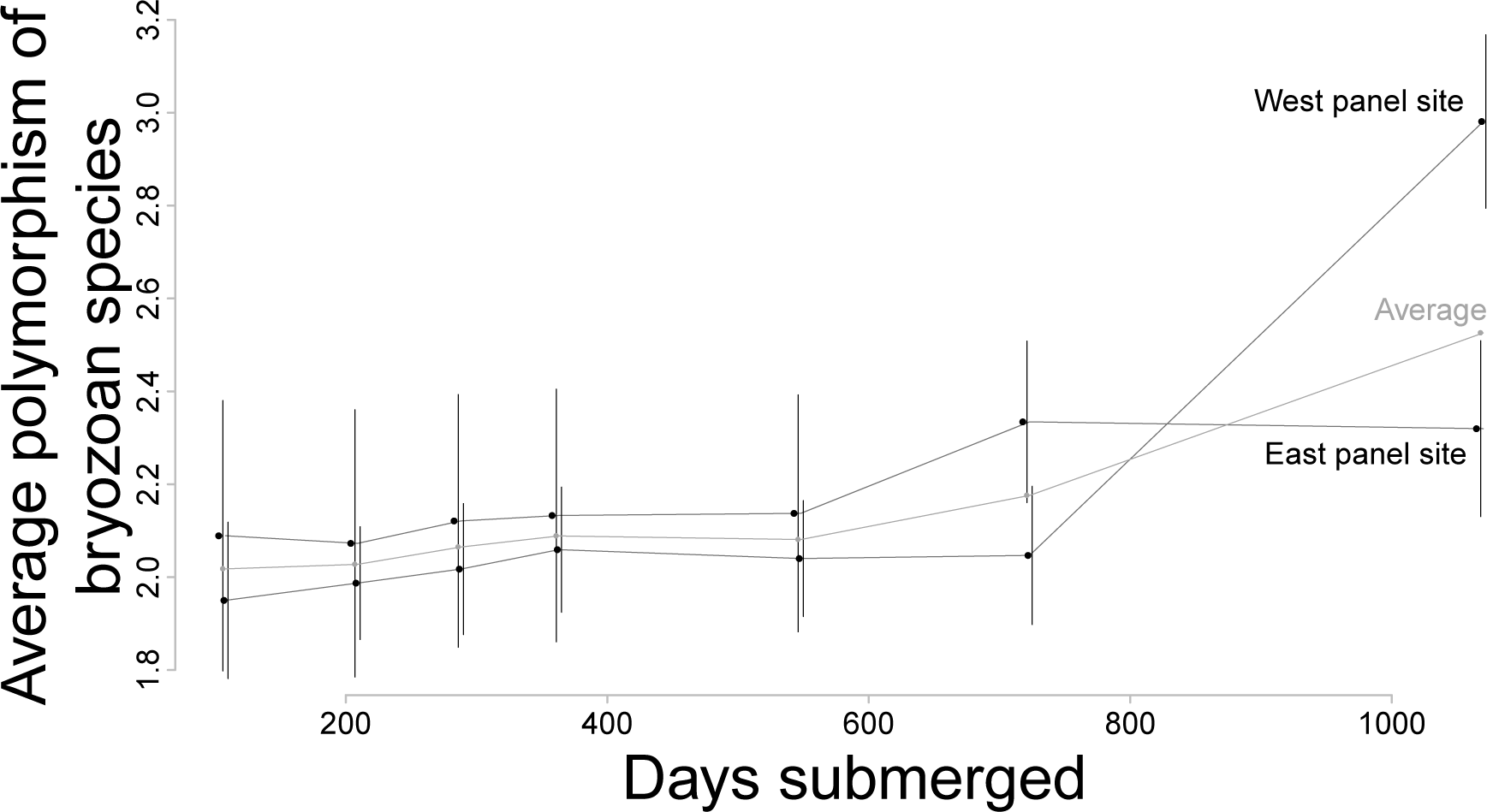
A direct comparison of competitive ability and polymorphism. Here we transform the percentage cover observed by Winston and Jackson (1984) to a relative proportion cover for each species to track the relative dominance of each species over time. We used the information in Fig. 3 to calculate the average number of polymorph types across species by weighting each species by its relative proportional cover. The average degree of polymorphism increases over three years despite the co-occurrence of species that differ in polymorphism. For both replicates pooled, the correlation of average polymorphism and time (Spearman’s *’)* is equal to 0.64 (P = 0.014). Time and polymorphism are also correlated in both replicate panels considered separately (west panel: Spearman’s *ρ* = 0.89, P = 0.012; east panel: Spearman’s *ρ* = 0.93, P = 0.008). The error bars summarize the variation among plates at each site and show one standard deviation on either side of the mean. We also offset the census days for east and west sites by 2 days to avoid overplotting of the points and error bars.

**Table 2:** Budding characteristics and polymorphism for 20 ascophoran cheilostome bryozoan species. The score is determined by the presence multizooidal budding, self-overgrowth, and frontal budding. Data on budding characteristics is from McKinney and Jackson (1991), Table 7.2.

| Species | Budding score | Polymorph types |
| --- | --- | --- |
| *Celleporella hyalina* | 1 | 1 |
| *Reptadeonella biparta* | 2 | 1 |
| *Escharella immersa* | 0 | 2 |
| *Fenestrulina malusi* | 0 | 2 |
| *Microporella cribosa* | 0 | 2 |
| *Reptadeonella costulata* | 2 | 2 |
| *Schizoporella errata* | 3 | 2 |
| *Schizoporella sanguinea* | 3 | 2 |
| *Schizoporella schizostoma* | 3 | 2 |
| *Schizoporella unicornis* | 3 | 2 |
| *Thalamoporella californica* | 1 | 2 |
| *Celleporella fusca* | 3 | 3 |
| *Escharina hynmanni* | 0 | 3 |
| *Microporella ciliata* | 0 | 3 |
| *Parasmittina raigii* | 3 | 3 |
| *Petraliella bisinuata* | 2 | 3 |
| *Rhynchozoon llarreyi* | 3 | 3 |
| *Schizoporella auriculata* | 3 | 4 |
| *Trematooecia aviculifera* | 3 | 4 |
| *Stylopoma spongites* | 3 | 6 |

## Discussion and Conclusion

The stability of the physical environment plays no direct role in the evolution of bryozoan polymorphism as postulated on the basis of environmental theory. In contrast, the incidence of polymorphism is strongly correlated with variations in ecological dominance and life-history as observed from both the increase in polymorphism generally as well as the increased dominance of exceptionally polymorphic species during ecological succession on the panels. This result is further borne out by an analysis of polymorphism of 20 ascophoran species ranked in terms of modes of budding that confer exceptional competitive ability and persistence in biological interactions (Table 7.2 in McKinney and Jackson 1991). The score is given by the sum of the presence or absence of multizooidal budding, self-overgrowth, and frontal budding. Only species with all three budding characteristics associated with dominance in competitive interactions have exceptionally high degrees of polymorphism (Table 2, Fig. 5). Conversely, species that lack one or more of these budding characteristic also lack diverse polymorph types (Fig. 5).

**Figure 5:**
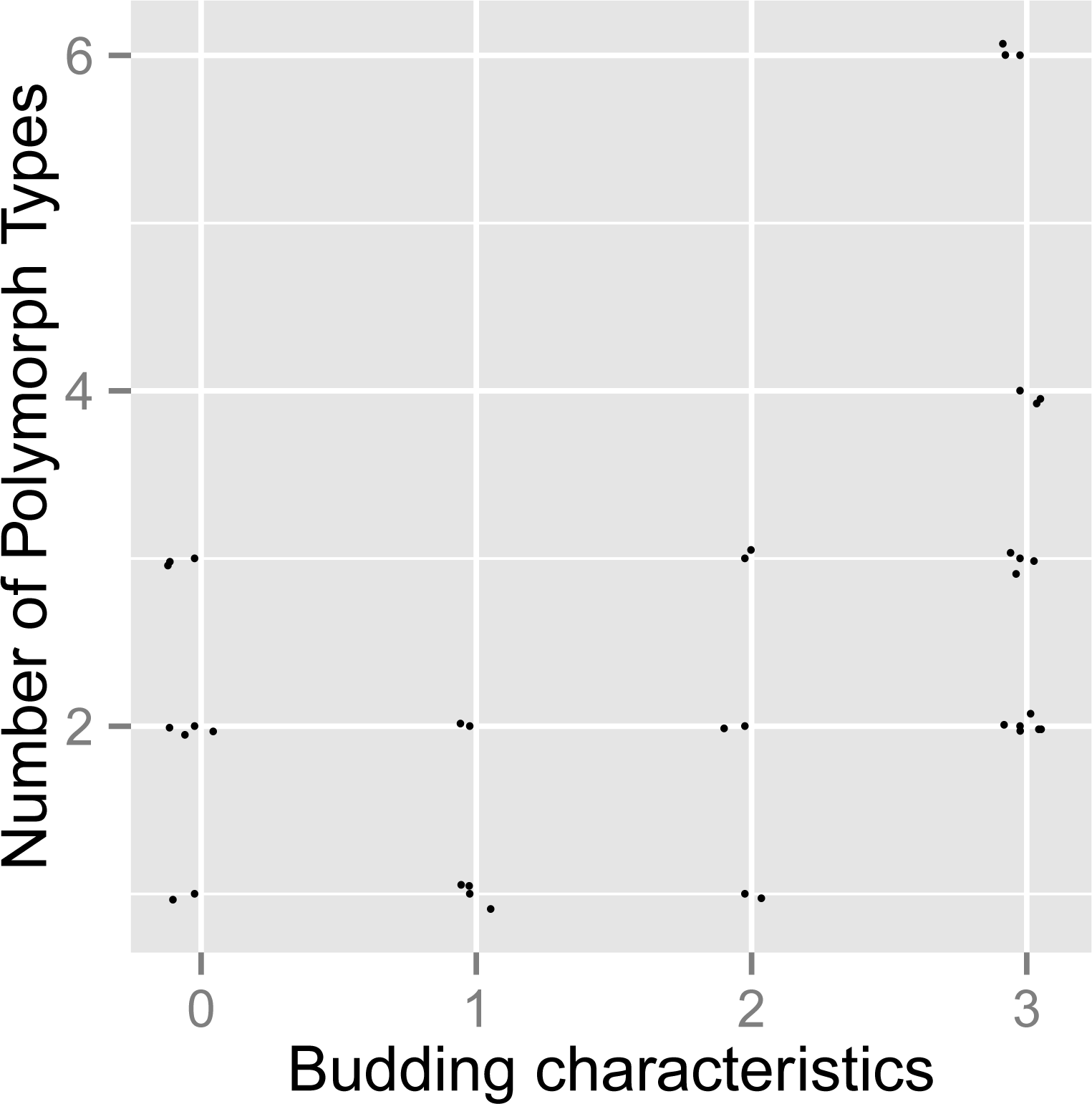
A comparison of budding characteristics and polymorphism of abundant encrusting ascophoran bryozoans. Bryozoans come from substrates that range in stability from algal fronds to corals and reef framework. Species are scored for the presence of three budding characteristics: multizooidal budding, self-overgrowth, and frontal budding. A species with all three budding characteristics will have a score of 3. Higher scores are correlated with an increased competitive ability. Data on budding characteristics is from McKinney and Jackson (1991), Table 7.2. and the species list, incidence of polymorphism and budding score used for this plot is shown in Table 2.

Opportunistic, early successional species tend to have lower levels of polymorphism than late successional species regardless of the overall oceanographic stability of their habitat. Rather than providing the time to accrue the benefits of expensive polymorphs, as expected by evolutionary theory, the stability of the coral habitats provides the time for ecological succession of different species to play out through a dense set of competitive ecological interactions. By specializing in particular life-history strategies, more bryozoan species, each with a different level of polymorphism, can coexist and interact.

Many rounds of succession can occur on the same substrate. Species with low levels of polymorphism are succeeded by species with higher levels so long as their substrate persists. However, physical disturbance or predation resulting in the death of an extremely polymorphic colony, such as many species of *Stylopoma* (Herrera et al. 1996; McKinney and Jackson 1991), permits the establishment of pioneering species with generally fewer polymorphic types. And so just as the death of a canopy tree in a tropical forest sets off a scramble among early successional species of trees (Connell 1978), the competitive cycle among bryozoans begins again.

How different types of modular polymorphs are developmentally specified still remains unknown. Within a single colony, all the diverse types of polymorphs are genetic clones. And so in some sense, the set of polymorphic types are each a part of a phenotypic reaction norm that the bryozoan genome is capable of. Yet the reaction norm perspective is an over simplification because of the discrete nature of modular polymorphic phenotypes and because not all types of polymorphs are derived from autozooids. Ovicells are derived from spines (Ostrovsky and Taylor 2004; Taylor and McKinney 2001) whereas avicularia are derived from autozooids (Banta 1973; Cheetham 1973). Thus, the modular form of a mother zooid may set limits on which types of polymorphs she and her descendants can produce.

There are hints that the evolution of polymorphism is linked to the evolution of life history strategies because species with more types of polymorphs tend to have less frequent reproductive specialists (Simpson 2012). Over macroevolutionary time there are two pathways to decrease that frequency that are contingent on the starting size of the most primitive colony. From originally small colonies, there must be a higher rate of increase in the number of “worker” or “soma” members relative to reproductives as the colony grows in size. Alternatively, when colonies are primitively large, colony members tend to loose reproductive competency even if the colony size remains largely the same.

In bryozoans, and perhaps other colonial and social animals, colonies with a single type of zooid will presumably have a life-history strategy that is tightly linked to the reproductive capabilities of its constituent zooids. If true, this implies that primitive colonies along both pathways inherit their life-histories from their members. From this perspective, understanding the macroevolution of polymorphism is critical to understanding the macroevolutionary proliferation of life history strategies.

